# A computational modelling framework to quantify the effects of passaging cell lines

**DOI:** 10.1101/161265

**Authors:** Wang Jin, Catherine J Penington, Scott W McCue, Matthew J Simpson

**Affiliations:** School of Mathematical Sciences, Queensland University of Technology (QUT) Brisbane, Queensland 4000, Australia

## Abstract

*In vitro* cell culture is routinely used to grow and supply a sufficiently large number of cells for various types of cell biology experiments. Previous experimental studies report that cell characteristics evolve as the passage number increases, and various cell lines can behave differently at high passage numbers. To provide insight into the putative mechanisms that might give rise to these differences, we perform *in silico* experiments using a random walk model to mimic the *in vitro* cell culture process. Our results show that it is possible for the average proliferation rate to either increase or decrease as the passaging process takes place, and this is due to a competition between the initial heterogeneity and the degree to which passaging damages the cells. We also simulate a suite of scratch assays with cells from near–homogeneous and heterogeneous cell lines, at both high and low passage numbers. Although it is common in the literature to report experimental results without disclosing the passage number, our results show that we obtain significantly different closure rates when performing *in silico* scratch assays using cells with different passage numbers. Therefore, we suggest that the passage number should always be reported to ensure that the experiment is as reproducible as possible. Furthermore, our modelling also suggests some avenues for further experimental examination that could be used to validate or refine our simulation results.

## Introduction

*In vitro* cell culture is routinely used to grow and supply cells for various types of cell biology experiments [1]. These experiments are used to study a wide range of biological phenomena including drug design, cancer spreading and tissue repair [9,20,23,42]. According to the American Type Culture Collection (ATCC) protocols, to grow cells in traditional two–dimensional (2D) *in vitro* cell culture, cells propagated in a growth medium are initially seeded as a monolayer in a cell culture flask [3], as shown in Fig 1a. Cells are seeded in a monolayer with a density typically varying from 10–20% of confluence [3]. Cells are then cultured in an incubator, in an appropriate temperature and CO_2_ concentration, and grown until they reach a density of 80%–90% of confluence [3]. To continue growing the population, cells are lifted, often using trypsin, and spilt into smaller proportions. The smaller subpopulations are transferred into new cell culture flasks to re-grow [3]. This process is referred to as *passaging*, with passage number indicating the number of splits [3, 4]. Although passaging is a standard process in 2D cell culture, the passage number of cells used in experiments is not always reported in experimental protocols [2, 17, 37–39, 41].

**Fig 1.**
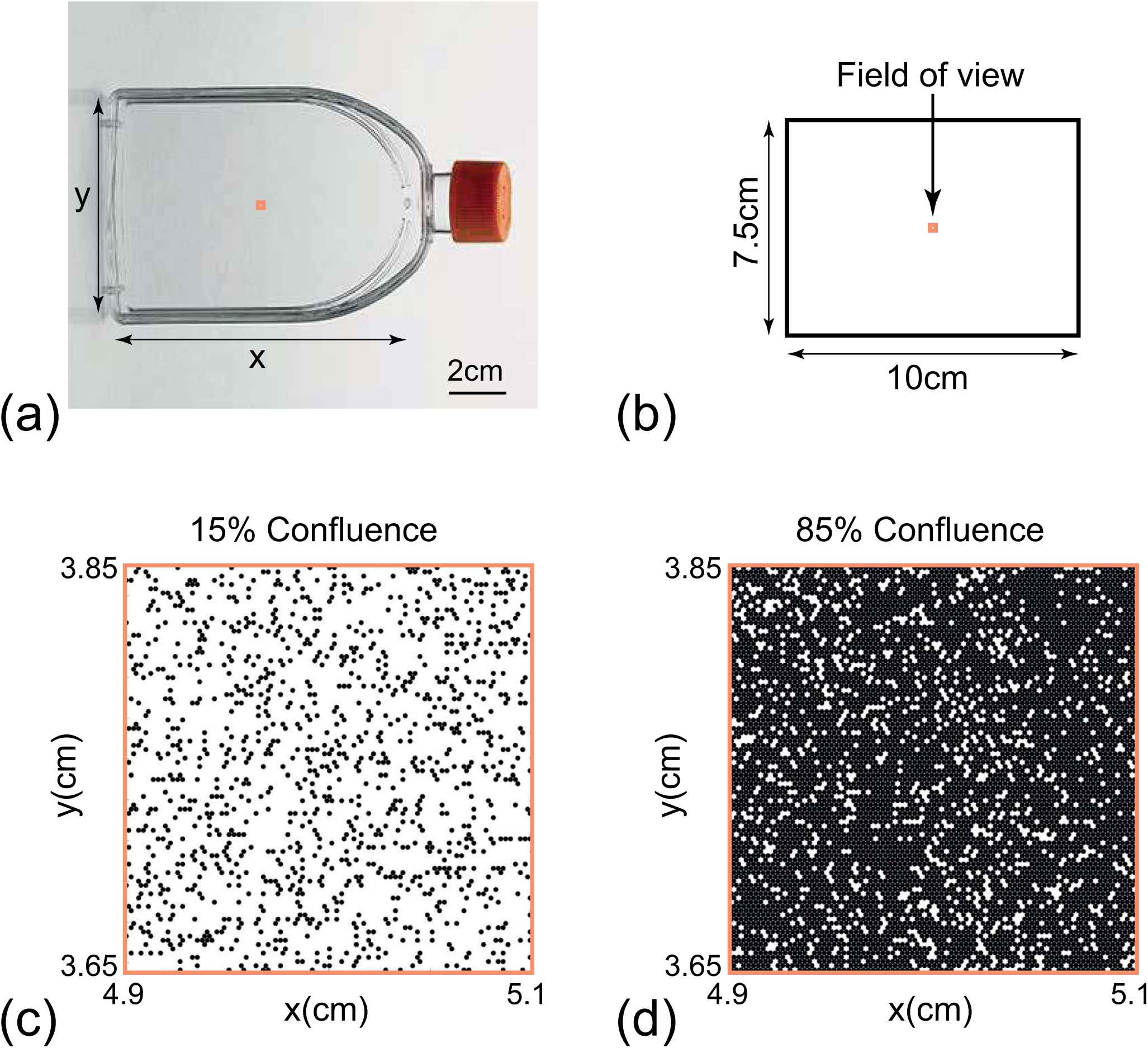
Schematic illustration of the simulation domain. (a) Photograph of a 75 cm^2^ cell culture flask. (b) Schematic of the 10 cm × 7.5 cm simulation domain that represents the 75 cm^2^ flask. The orange squares in (a) and (b) indicate the 2 mm × 2 mm field of view. (c) Snapshot of the field of view at 15% confluence. (d) Snapshot of the field of view at 85% confluence.

It is known that passaging can affect cells in a number of ways, and therefore has the potential to impact the reproducibility of *in vitro* experiments [41]. There are many ways in which passaging can affect cells. For example, primary cells, which are directly isolated from living tissues [16], undergo morphological changes and cumulative damage as the passage number increases [6, 10, 18, 22, 24, 31, 32, 34]. As a result, the cell morphology, migration rate and proliferation rate can become increasingly varied, which is thought to increase the heterogeneity in cell lines [10, 18, 24, 32, 34]. Because a range of cell behaviours could depend on passage number, the passaging process can be a source of variability that affects the reproducibility of various *in vitro* experiments, such as 2D scratch assays [2, 4, 41].

Seemingly contradictory observations have been reported about the effects of passaging cell lines [10, 18, 26, 32, 34]. For example, Hayflick reports that for human diploid cell lines, cells at high passage numbers demonstrate increased generation time, gradual cessation of mitotic activities, and accumulation of cellular debris [18]. This observation of decreased cell proliferation rate is also supported by studies of other cell lines [10, 32, 34]. However, Lin and coworkers show that the population of LNCaP cells at passage number 70 is over two times larger than that at passage number 38 after five days [26]. It has also been stated that for some cell lines, changes due to the passaging process occur at relatively low passage numbers, whereas for other cell lines the changes occur at relatively high passage numbers [4]. Therefore, we are motivated to undertake a mechanistic study to quantify how different variables relevant to the passaging process might give rise to such seemingly contradictory observations and to explore how these effects might impact the reproducibility of *in vitro* experiments.

Although problems associated with high passage numbers are widely acknowledged, the mechanism of passage–induced changes is not well understood [4, 10, 13, 18, 26, 29, 32, 34, 45]. For example, standard experimental protocols suggest avoiding cells at high passage numbers, whereas the definition of a ‘high passage number’ is rather vague [4, 29]. On the other hand, the mechanism that causes the seemingly contradictory observations at high passage numbers still remains unknown [10, 18, 26, 32, 34]. Computational models can be useful for exploring mechanisms and trade-offs between various factors. Therefore, the problems with high passage numbers invoke us to apply a computational model to investigate putative mechanisms that could lead to the seemingly contradictory changes. As far as we are aware, this is the first time that problems with passaging of cell lines are investigated using a computational approach of this kind.

In this work, we describe a mathematical model that can be used to study the passaging process in 2D *in vitro* cell culture [19, 43]. A key feature of our model is that we allow individual cells within the population to take on a range of characteristics, such as variable proliferation rates, and therefore it is natural to focus on using a discrete model for this purpose [8, 19]. In particular, we are interested in examining whether the apparently contradictory effects of passaging reported in the literature can be recapitulated using a fairly straightforward discrete model. After examining the trade-off between cell heterogeneity and passage–induced damage, we then use the *in silico* model to examine how the passaging process might affect the reproducibility of scratch assays [20, 25]. In this work we focus on the impact of passaging on the cell proliferation rate, and apply a discrete model to explain how passaging can lead to either increasing or decreasing proliferation rates, depending on the competing effects of natural inheritance versus passaging–induced damage. In our model we impose three key assumptions: (i) the passaging process does not affect the cells’ ability to migrate; (ii) initially the proliferation rate of each cell is assigned randomly from a normal distribution; and (iii) when proliferating, daughter cells inherit the same proliferation rate as the mother cell. Our approach is to focus on two prototype cell populations. The first is near–homogeneous in the sense that the proliferation rate of the cells is close to constant throughout the population initially. The second has a distinctively heterogeneous distribution of proliferation rates. For each prototype population, we systematically vary the amount of damage caused by passaging to investigate the impact of the damage.

## Discrete model

### Model framework

We use a discrete random walk model to simulate the passaging process and we refer to individual random walkers in the model as cells. All simulations are performed on a hexagonal lattice, with the lattice spacing Δ taken to be equal to the average cell diameter [19]. The model includes crowding effects by ensuring that there is, at most, one cell per lattice site [36]. Each lattice site, indexed (*i, j*) where *i, j ∈* ℤ^+^, has position

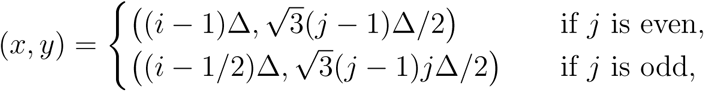

such that 1 ≤ *i* ≤ *I* and 1 ≤ *j* ≤ *J* [19]. In any single realisation of the model, the occupancy of site (*i, j*) is denoted *C*_*i,j*_, with *C*_*i,j*_ = 1 if the site is occupied, and *C*_*i,j*_ = 0 if vacant.

If there are *N* (*t*) cells at time *t*, then during the next time step of duration *τ, N* (*t*) cells are selected independently at random, one at a time with replacement, and given the opportunity to move [19, 36]. The randomly selected cell attempts to move, with probability *P*_*m*_, to one of the six nearest neighbour sites, with the target site chosen randomly. Motility events are aborted if a cell attempts to move to an occupied site. Once *N* (*t*) potential motility events are attempted, another *N* (*t*) cells are selected independently, at random, one at a time with replacement, and given the opportunity to proliferate with probability *P*_*p*_. The location of the daughter cell is chosen, at random, from one of the six nearest neighbour lattice sites [19, 36]. If the selected lattice site is occupied, the potential proliferation event is aborted. In contrast, if the selected site is vacant, a new daughter cell is placed on that site. After the *N* (*t*) potential proliferation events have been attempted, *N* (*t* + *τ*) is updated [19, 36].

The discrete models in this study are coded in C++. And the C++ simulation code is supplied in the Supporting Information.

### Simulation domain

The domain is a rectangle of dimensions 10 cm by 7.5 cm, which we use to represent the 75 cm^2^ cell culture flask in Fig 1a. This corresponds to a simulation domain in which *I* = 4168 and *J* = 3610, with Δ = 24 *μ*m [20]. Therefore, the maximum number of cells in a 100% confluent monolayer is approximately 15 million. To simplify our visualisation of the model output, although we always perform simulations on the entire 10 cm by 7.5 cm simulation domain, we visualise a smaller, 2 mm by 2 mm, subregion in the centre of the simulation domain, as shown in Fig 1b. No flux boundary conditions along the boundaries of the simulation domain are applied in all cases. For the remainder of this work, we visualise snapshots of the distribution of cells in the smaller field of view, such as the results in Fig 1c–d.

### Initial condition

Simulations are initiated by randomly populating 15% of lattice sites [3]. At each passage number, the growth of cells in the culture is terminated when 85% confluence is reached. The migration probability *P*_*m*_ of each cell is held constant. Motivated by experimental data of the duration of the mitotic phase for individual cells [15], each cell is initially assigned a random value of *P*_*p*_, drawn from a normal distribution *𝒩* (*μ*_*p*_, *σ*) to mimic the stochasticity in proliferation rate among the initial population. When a proliferation event takes place, we invoke the simplest mechanism by assuming that both daughter cells inherit the proliferation rate of the mother cell. For all simulations we set *P*_*m*_ = 0.35, *μ*_*p*_ = 0.004, Δ = 24 *μ*m and *τ* = 1/12 h so that we are considering cell populations with a typical cell diameter, cell diffusivity (*D* ≈ 600 *μ*m^2^/h) and average proliferation rate (*λ* ≈ 0.05 /h) [19, 36]. We consider two prototype cell populations: (i) a near–homogeneous cell population with a relatively small variance, *σ* = 1 × 10^−4^; and (ii) a heterogeneous cell population with a larger variance, *σ* = 1 × 10^−3^. We choose the values of the standard deviation *σ*, so that the proliferation rate distribution is within a biologically reasonable range, and the degree of heterogeneity in the near–homogeneous and heterogeneous cell lines are distinguishable.

### Passaging

In our simulations passaging takes place immediately after the population grows to 85% confluence [3]. To split the populations we randomly select a number of cells that is equivalent to cover 15% of lattice sites. These cells are randomly placed on an empty simulation domain to mimic the splitting of cells in the passaging process. Note that *P*_*m*_ is constant for all cells whereas we allow *P*_*p*_ to vary amongst the population and we also assume that the process of passaging the cells involves some damage [32]. Considering that the passaging process involves a combination of chemical (e.g. the usage of trypsin) and mechanical disturbances known to disrupt normal cell behavior, it is reasonable to incorporate some kind of damage mechanism into the passage simulations [3, 6, 10, 18, 31, 32, 34]. However, the exact cause and the form of the passage–induced damage have not been established. Therefore any form of the passage–induced damage which illustrates certain degrees of stochasticity could be a reasonable choice. In this study, we consider two different degrees of passage–induced damage:

- Small amount of damage: *P*_*p*_ of each cell is decreased by ∊, where ∊∼ *𝒩* (2 × 10^−5^, 2 × 10^−5^);
- Large amount of damage: *P*_*p*_ of each cell is decreased by ∊, where ∊∼ *𝒩* (1 × 10^−4^, 1 × 10^−4^).

Each time the population of cells is split, the passage number increases by one. As previous studies indicate that cell proliferation increases at high passage numbers [26], it is possible to assume that the passage–induced damage could lead to the increase in proliferation rate. However, since the aim of this study is to examine the trade-offs between the initial heterogeneity in cell proliferation and the passage–induced damage, in both scenarios we only consider non– negative passage–induced damage by changing any negative damage to zero. This assumption allows us to limit the factors that can increase cell proliferation.

## Results and discussion

### Passaging cell lines without passage–induced damage

We first investigate how the initial degree of heterogeneity in proliferation rate changes as the passage number increases. In this first set of results we do not consider any form of passage–induced damage. We consider a suite of simulations from passage number 0 to 30 and present results for both the near–homogeneous cell line and the heterogeneous cell line. Snapshots of the field of view at passage number 0 and passage number 30, for both prototype cell populations, are shown in Fig 2a–d and Fig 3a–d, respectively. In each snapshot, different colours of cells represent different ranges of the proliferation rate, with red indicating the fastest–proliferating cells and blue showing the slowest–proliferating cells. At the end of passage number 0 we observe a larger variation in cell proliferation rate in the heterogeneous cell line than the near–homogeneous cell line, as we might expect. At the end of passage number 30 we see that there is a dramatic change in the average proliferation rate of cells in the heterogeneous cell line. This change is caused by the fact that cells with higher proliferation rates are more likely to produce daughter cells that directly inherit the higher proliferation rate of the mother cell. Therefore, we observe a greater proportion of faster–proliferating cells in the heterogeneous cell line at high passage number. This leads to a larger average value of *P*_*p*_ and a greater variation in *P*_*p*_ across the whole population of cells in the heterogeneous cells line, as shown in Fig 2e–h and Fig 3e–h, respectively.

**Fig 2.**
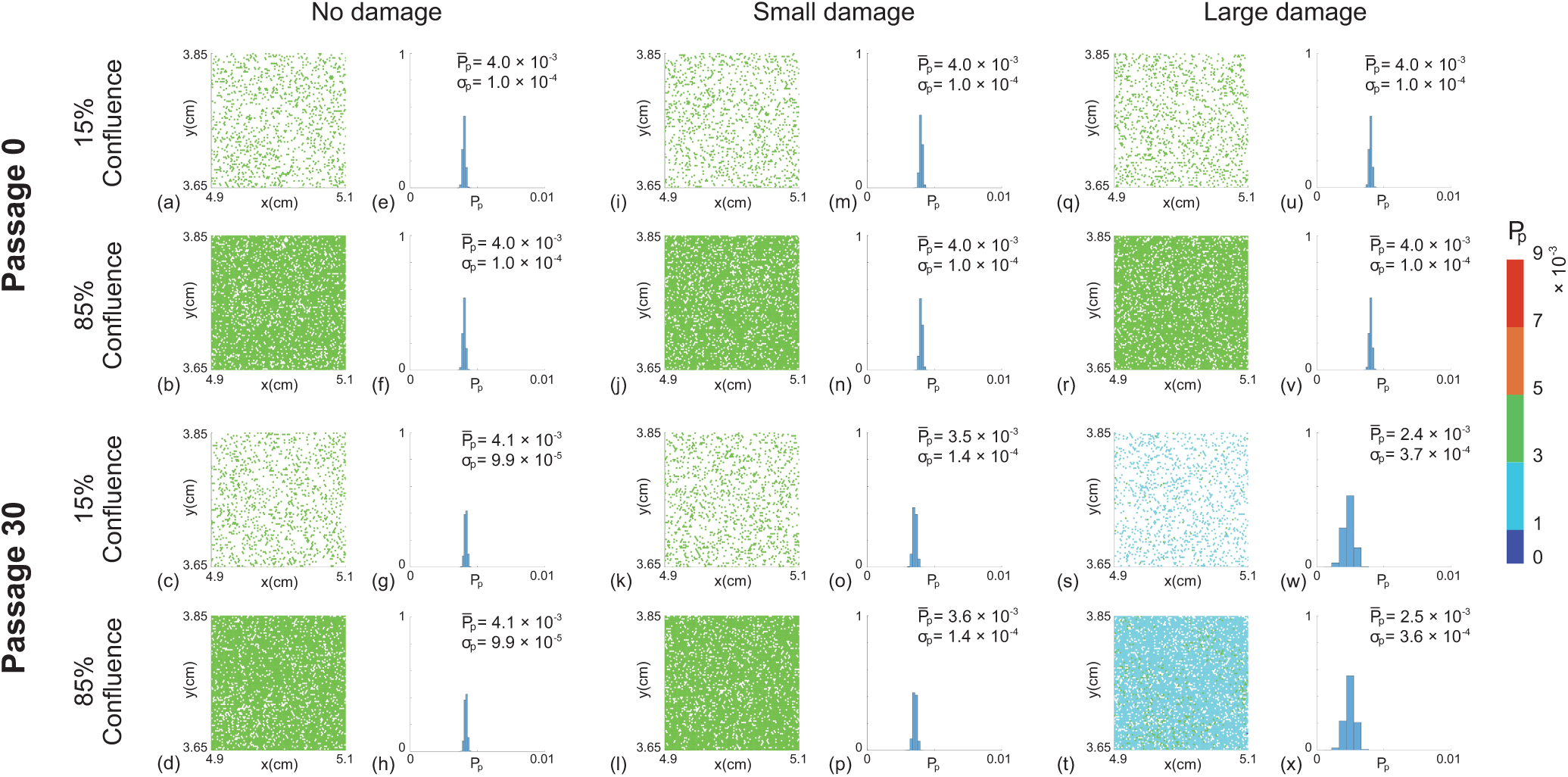
Snapshots of simulations for a near–homogeneous cell line. For each passage number, snapshots at the beginning (15% confluence) and end (85% confluence) of the experiments are shown. Results in (a)–(d),(i)–(l) and (q)–(t) show snapshots of the field of view at passage number 0 and 30, with ∊ = 0, ∊ ∼ *𝒩* (2 × 10^−5^, 2 × 10^−5^) and ∊ ∼ *𝒩* (1 × 10^−4^, 1 × 10^−4^), respectively. Results in (e)–(h),(m)–(p) and (u)–(x) show distributions of *P*_*p*_ for the entire domain at passage number 0 and 30, with ∊ = 0, ∊ ∼ *𝒩* (2 × 10^−5^, 2 × 10^−5^) and ∊ ∼ *𝒩* (1 × 10^−4^, 1 × 10^−4^), respectively. The distribution of *P*_*p*_ in each subfigure is obtained from one single realisation. The colour bar indicates *P*_*p*_ for individual cells. 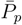 and *σ*_*p*_ represent the mean and standard deviation of *P*_*p*_. Each simulation is initiated by randomly populating 15% of lattice sites, on a lattice of size *I* = 4168 and *J* = 3610, with *P*_*p*_ ∼ *𝒩* (4 × 10^−3^, 1 × 10^−4^) for each cell. All simulations correspond to Δ = 24 *μ*m, *τ* = 1/12 h, and *P*_*m*_ = 0.35.

**Fig 3.**
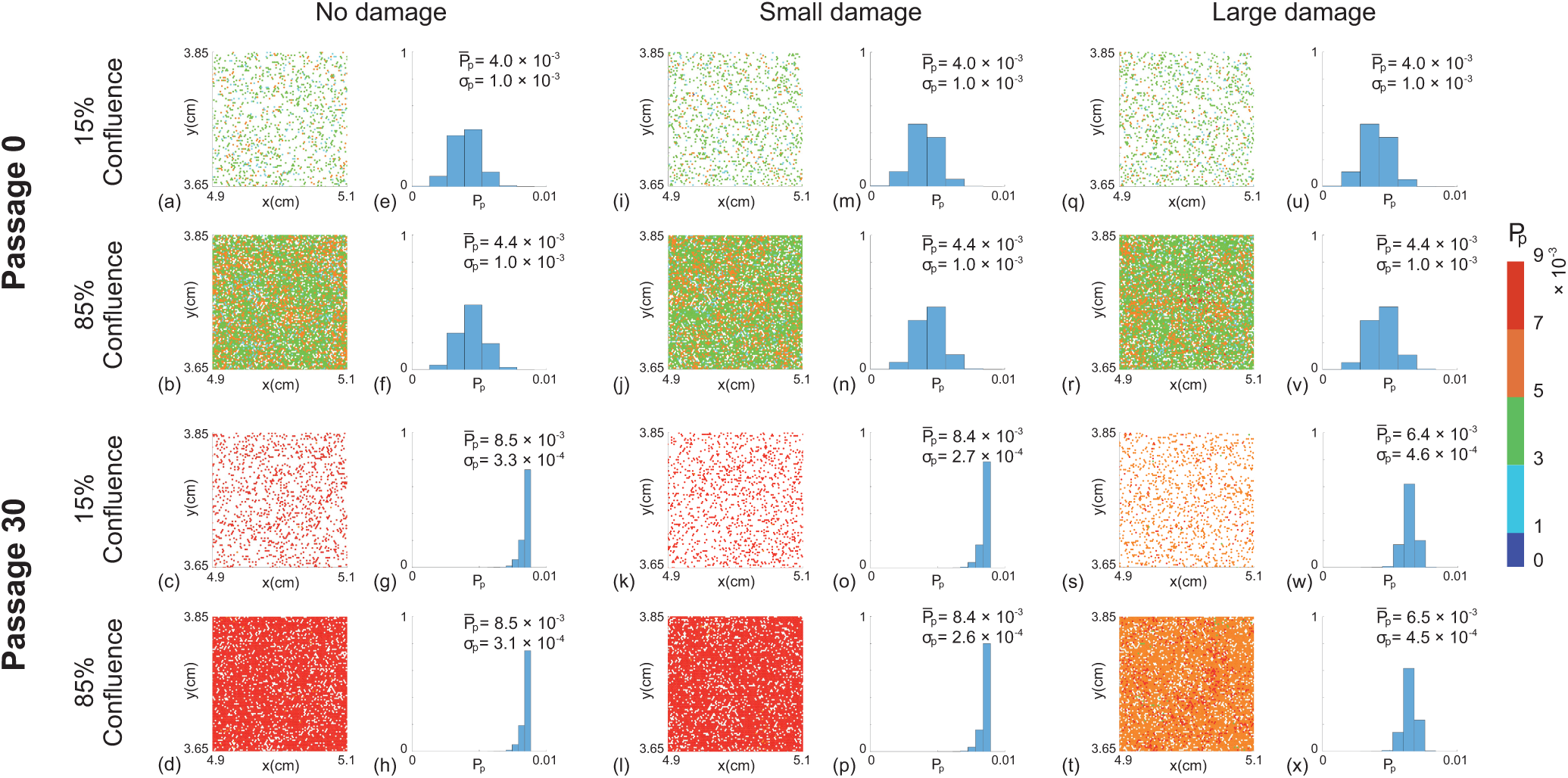
Snapshots of simulations for a heterogeneous cell line. For each passage number, snapshots at the beginning (15% confluence) and end (85% confluence) of the experiments are shown. Results in (a)–(d),(i)–(l) and (q)–(t) show snapshots of the field of view at passage number 0 and 30, with ∊ = 0, ∊ ∼ *𝒩* (2 × 10^−5^, 2 × 10^−5^) and ∊ ∼ *𝒩* (1 × 10^−4^, 1 × 10^−4^), respectively. Results in (e)–(h),(m)–(p) and (u)–(x) show distributions of *P*_*p*_ for the entire domain at passage number 0 and 30, with ∊ = 0, ∊ ∼ *𝒩* (2 × 10^−5^, 2 × 10^−5^) and ∊ ∼ *𝒩* (1 × 10^−4^, 1 × 10^−4^), respectively. The distribution of *P*_*p*_ in each subfigure is obtained from one single realisation. The colour bar indicates *P*_*p*_ for individual cells. 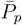 and *σ*_*p*_ represent the mean and standard deviation of *P*_*p*_. Each simulation is initiated by randomly populating 15% of lattice sites, on a lattice of size *I* = 4168 and *J* = 3610, with *P*_*p*_ ∼ *𝒩* (4 × 10^−3^, 1 × 10^−3^) for each cell. All simulations correspond to Δ = 24 *μ*m, *τ* = 1/12 h, and *P*_*m*_ = 0.35.

To summarise how the cell proliferation rate changes with passage number, we plot the evolution of the proliferation rate data from the entire populations as gboxplots [28] in Fig 4. The boxplots show the median and quartiles of the distribution of *P*_*p*_ from the entire population as a function of the passage number. Comparing results in Fig 4a and Fig 4d shows that the median *P*_*p*_ increases much faster in the heterogeneous cell line than the near–homogeneous cell line. For the near–homogeneous cell line the distribution of *P*_*p*_ appears to be approximately independent of the passage number in this case. In contrast, the distribution of *P*_*p*_ for the heterogeneous cell line is strongly dependent on the passage number. In particular, the median *P*_*p*_ increases, and the distribution of *P*_*p*_ becomes increasingly negatively skewed as the passage number increases. Overall, these results suggest that starting with the same average proliferate rate, the degree of heterogeneity of the cell line alone is enough to lead to very different outcomes when the two cell lines are sufficiently passaged. Therefore, the initial heterogeneity of the cell line appears to be important in terms of understanding how passaging affects properties of cell lines.

**Fig 4.**
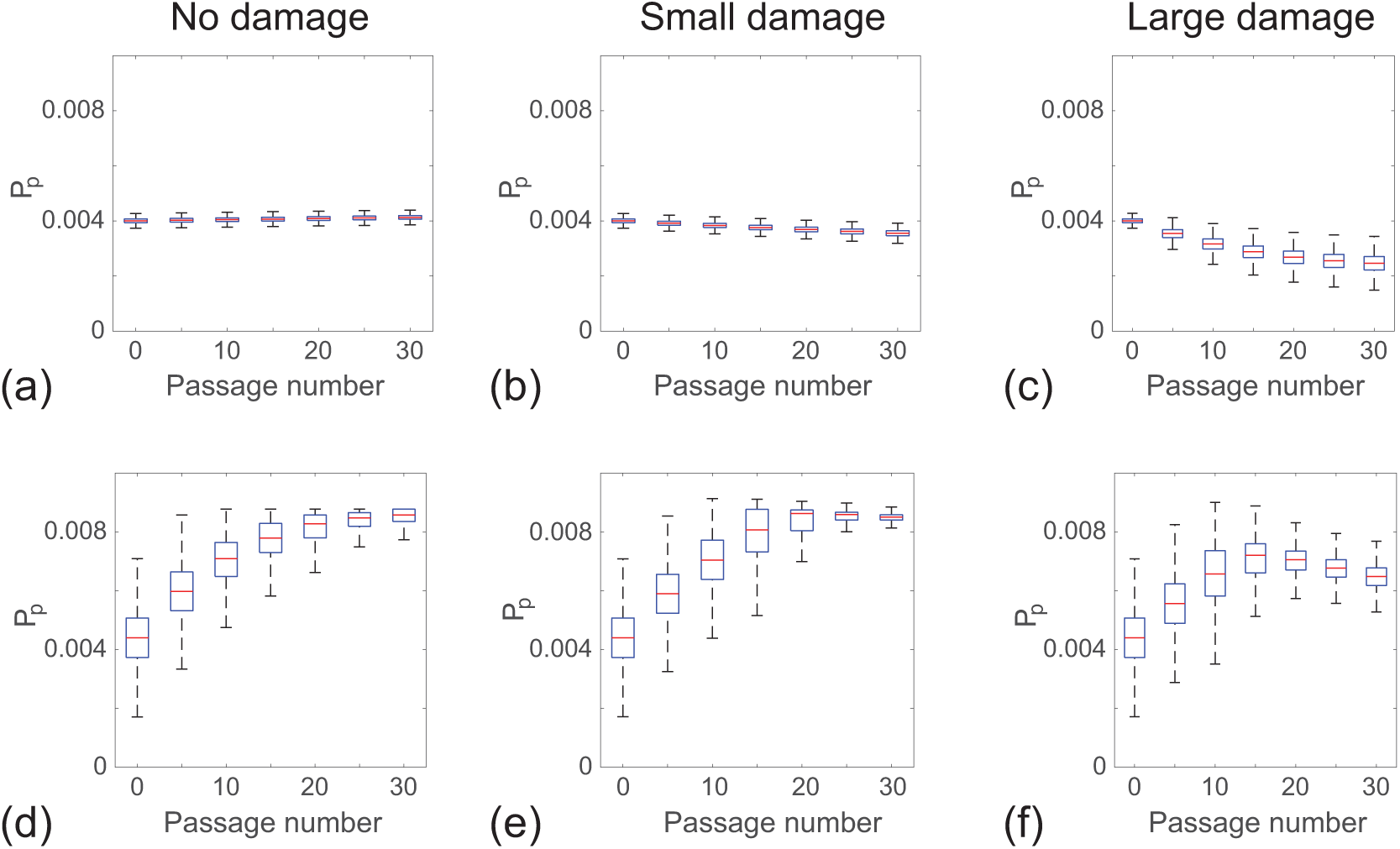
Distribution of *P*_*p*_ as a function of passage number. Results in (a)–(c) show boxplots of *P*_*p*_ for a near–homogeneous cell line at 85% confluence for: (a) no damage ∊ = 0; (b) small amount of damage, ∊ ∼ *𝒩* (2 × 10^−5^, 2 × 10^−5^); and (c) large amount of damage, ∊ ∼ *𝒩* (1 × 10^−4^, 1 × 10^−4^). Results in (d)–(f) show boxplots of *P*_*p*_ for a heterogeneous cell line at 85% confluence for: (d) no damage ∊ = 0; (e) small amount of damage, ∊ ∼ *𝒩* (2 × 10^−5^, 2 × 10^−5^); and (f) large amount of damage, *E ∼ 𝒩* (1 × 10^−4^, 1 × 10^−4^). In each subfigure the distribution of *P*_*p*_ at individual passage numbers is obtained from one single realisation. Each simulation is initiated by randomly populating 15% of lattice sites on a lattice of size *I* = 4168 and *J* = 3610, with *P*_*p*_ ∼ *𝒩* (4 × 10^−3^, 1 × 10^−4^) for the near–homogeneous cell line and *P*_*p*_ ∼ *𝒩* (4 × 10^−3^, 1 × 10^−3^) for the heterogeneous cell line. All simulations correspond to Δ = 24 *μ*m, *τ* = 1/12 h, and *P*_*m*_ = 0.35.

In this first set of results, we find that differences in the cell proliferation rate among the cell population can lead to changes in the overall population behaviour at sufficiently high passage numbers. We note that in both prototype cell populations, the average proliferation rate increases with the passage number and this is consistent with some previous experimental studies [26]. However, most experimental studies report a decrease in average proliferation rate with increasing passage number [10, 18, 32, 34]. This observation motivates us to include a second mechanism in our discrete model, namely passage–induced damage.

### Passaging cell lines with passage–induced damage

We now investigate the impact of including passage–induced damage, and we consider both small and large amounts of damage scenarios. All other features of our simulations are maintained as described in the section without passage–induced damage. Snapshots of simulations including small and large amounts of damage, and boxplots showing the distribution of *P*_*p*_ data are shown in Fig 2–4. Comparing results in Fig 2a-h and Fig 2i-p suggests that we observe very similar outcomes when we include a small amount of damage in the simulations of the near–homogeneous cell line. Similarly, results in Fig 3a-h and Fig 3i-p suggest that the small amount of damage has a negligible impact on the passaging process for the heterogenous cell line. In contrast, with the large amount of damage we see that the proliferation rate decreases by passage number 30 in the near–homogeneous cell line, as shown in Fig 2q–x, whereas results in Fig 3q–x show that the proliferation rate increases by passage number 30, but the increase in proliferation rate is not as pronounced as in the case where there is no damage in the heterogenous cell line.

Results in Fig 2–3 focus on snapshots of the population at passage numbers 0 and 30. Additional results in Fig 4b–c and Fig 4e–f to show how the distribution of *P*_*p*_ evolves as a function of the passage number. For the near–homogeneous cell line, the median *P*_*p*_ decreases monotonically with the passage number for both small and large amounts of damage. In contrast, the median *P*_*p*_ for the heterogeneous cell line behaves very differently as it increases until approximately passage number 20, and then decreases with further passaging. These results, combined, provide a simple explanation for why some previous studies have reported that the proliferation rate can increase with passage number, as in the case of Fig 4d–e, whereas other studies suggest that the proliferation rate can decrease with passage number, as in the case of Fig 4c. In fact, our results suggest that it is possible to have a situation where the proliferation rate both increases and decreases with passage number, as in the case of Fig 4f, and we observe different trends depending on the passage number. These differences arise in our model due to a trade-off between the initial heterogeneity of the cell line and the amount of damage sustained in the passaging process.

### Scratch assay with passaged cells

Having demonstrated that the interplay between cell heterogeneity and passage–induced damage can lead to complicated trends in the relationship between the proliferation rate and passage number, it is still unclear how these kinds of differences can affect how we interpret *in vitro* experiments. To explore this issue we use cells from a range of passage conditions to mimic a scratch assay [25]. For this purpose we focus on the geometry associated with experimental images obtained from an IncuCyte ZOOM™ scratch assay [11, 14, 20, 33, 44], as shown in Fig 5. The images, of dimension 1400 *μ*m × 1900 *μ*m, show a fixed field of view that is much smaller than the spatial extent of the cells in the scratch assay [19, 21]. To model this situation we apply zero net flux boundary conditions along all boundaries of the lattice. We use a lattice of size 80 × 68 to accommodate a typical population of cells with Δ = 24 *μ*m. To initialise the scratch assay, we randomly populate all lattice sites with an equal probability of 30% [20]. To simulate the scratch, we remove all cells from a vertical strip of width approximately 550 *μ*m, and we then observe the rate at which the populations spread into the vacant area. All cells have the same constant value of *P*_*m*_ = 0.35, and we assign values of *P*_*p*_ by sampling from the various histograms in Fig 2 and 3. This means that we are effectively simulating a scratch assay using cells from different cell lines, with different amounts of passage-induced damage, and from different passage numbers according to our *in silico* results of cell culture growth in the previous section.

**Fig 5.**
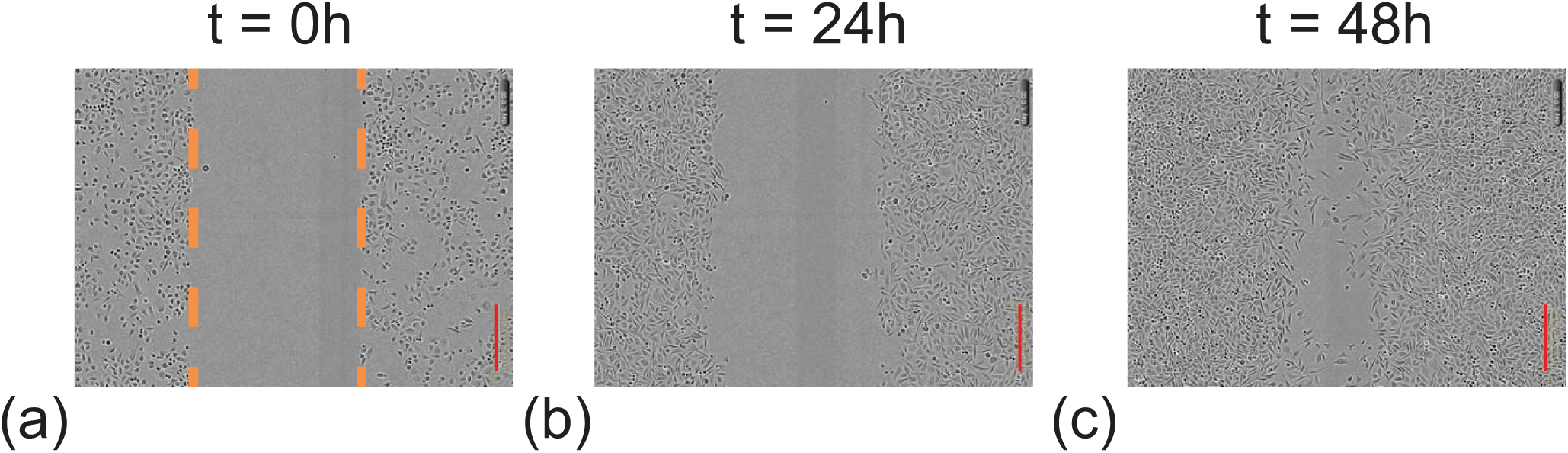
Experimental images of IncuCyte ZOOM™ scratch assay [20, 21]. The images in (a)–(c) show the closure of the initially scratched region which is highlighted by the dashed orange lines at *t* = 0. The red scale bar corresponds to 300 *μ*m.

Snapshots from the discrete model, showing the progression of the scratch assays, are shown in Fig 6–7. In general we see that, regardless of the initial cell population, all of the scratch assays lead to successful closure by approximately 48-72 h, which is consistent with standard experimental observations [14, 20]. However, close examination of the results reveals some differences. In particular, visual inspection of the snapshots suggests that those cell populations with higher initial proliferation rate lead to larger numbers of cells at later times, and hence more rapid closure of the initially–vacant space. These trends are subtle, but are most obvious in Fig 7 where the population corresponds to cells taken from passage number 30, with no damage, leading to more effective re-colonisation of the initially–vacant space than cells from passage number 0. Since these differences are subtle it may be difficult to detect them when visually comparing results from scratch assays. Therefore, we will now quantify the spatial and temporal distribution of cells in Fig 6–7 to provide more information.

**Fig 6.**
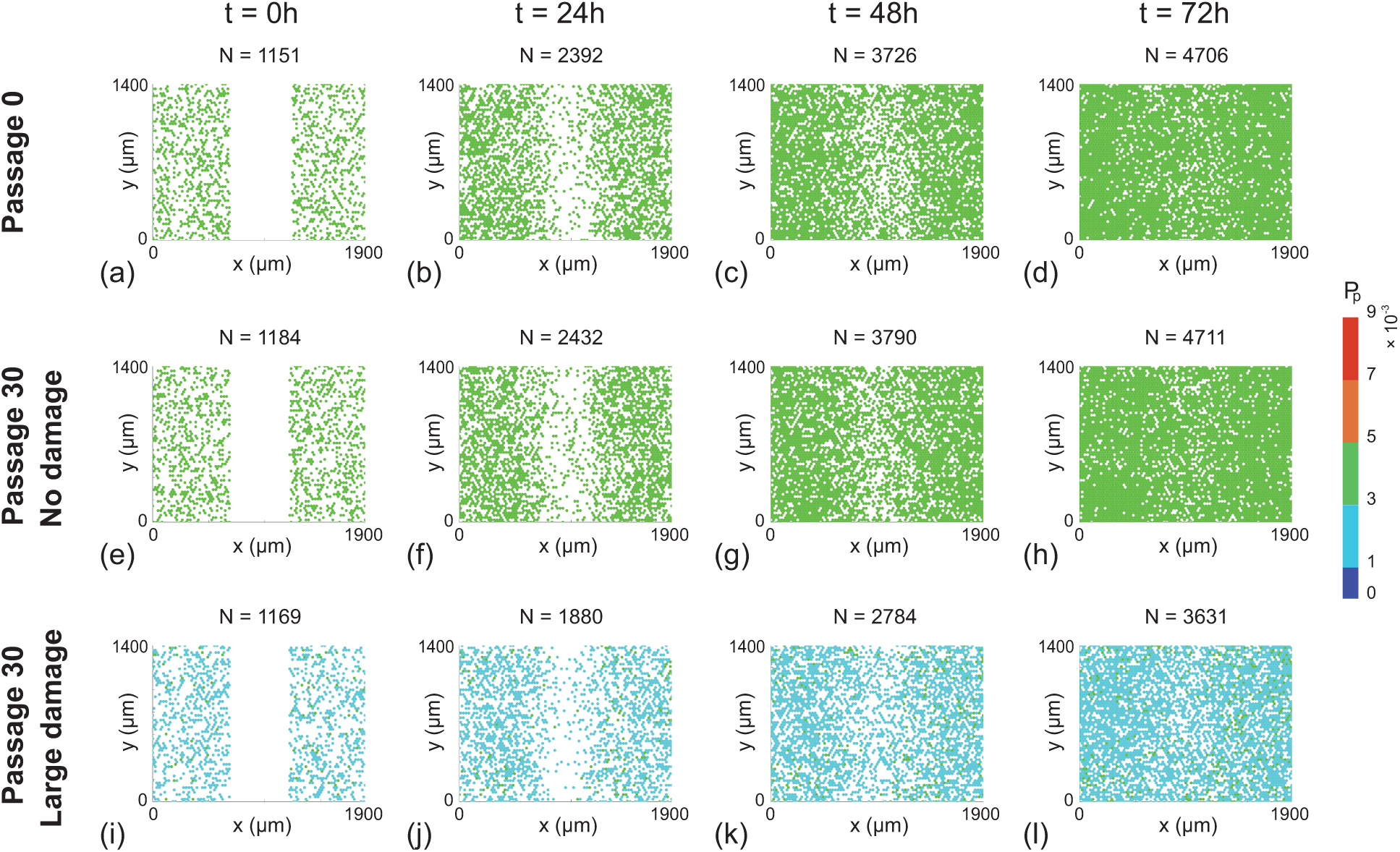
Snapshots of a suite of scratch assays performed using a near–homogeneous cell line. In each column the distributions of cells at time *t* = 0, 24, 48, 72 h are shown. Each simulation is initiated by randomly populating a lattice of size 80 × 68, so that each site is occupied with probability 30%. A scratch of 23 lattice sites wide is made at *t* = 0 h. All simulations correspond to Δ = 24 *μ*m, *τ* = 1/12 h, and *P*_*m*_ = 0.35. In each row the initial *P*_*p*_ of individual cells is assigned by randomly selecting from the *in silico* data in Fig 2f, h and x, respectively.

**Fig 7.**
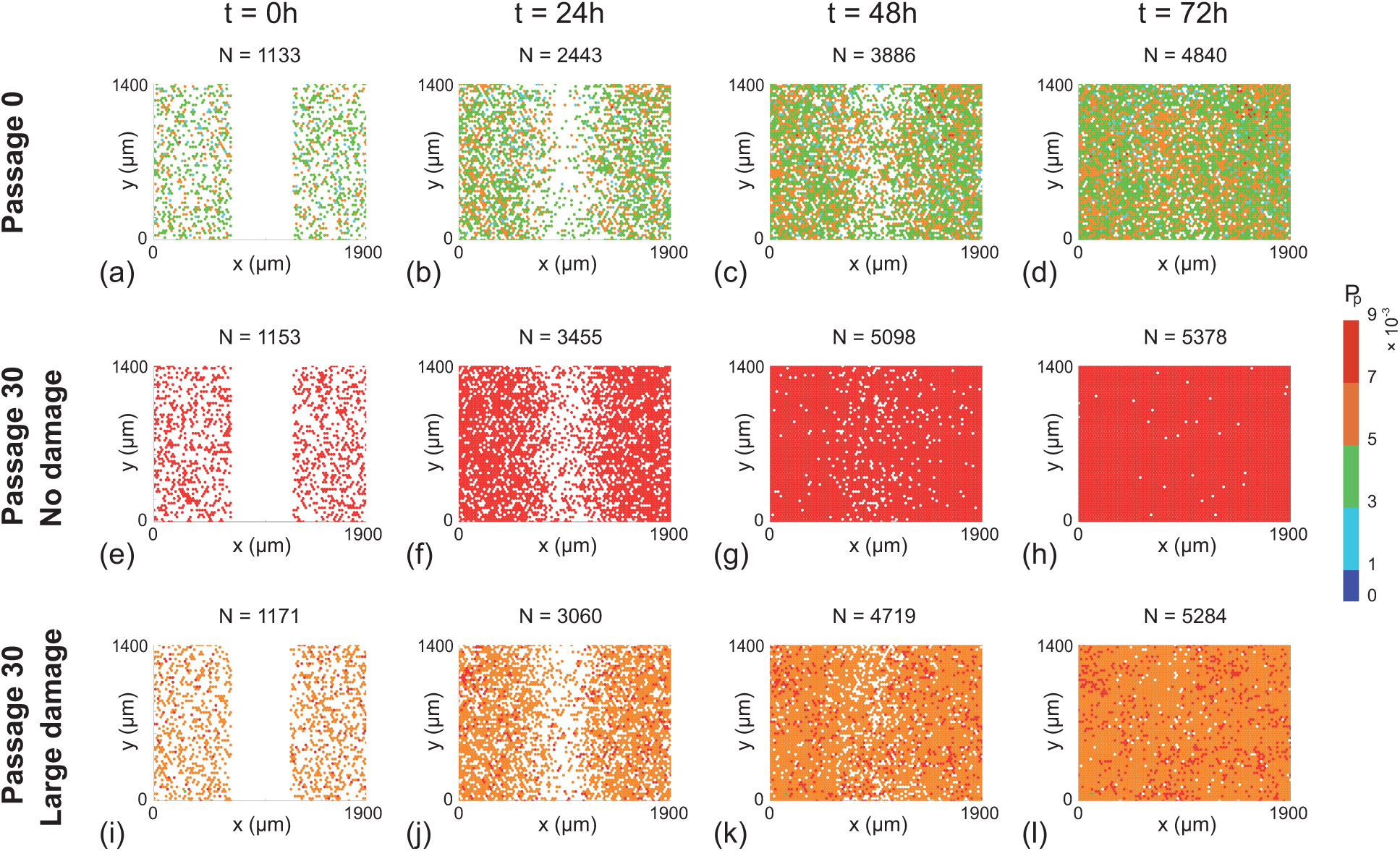
Snapshots of a suite of scratch assays performed using a heterogeneous cell line. In each column the distributions of cells at time *t* = 0, 24, 48, 72 h are shown. Each simulation is initiated by randomly populating a lattice of size 80 × 68, so that each site is occupied with probability 30%. A scratch of 23 lattice sites wide is made at *t* = 0 h. All simulations correspond to Δ = 24 *μ*m,*τ* = 1/12 h, and *P*_*m*_ = 0.35. In each row the initial *P*_*p*_ of individual cells is assigned by randomly selecting from the *in silico* data in Fig 3f, h and x, respectively.

Since the initial condition is uniform in the vertical direction [19, 36], we average the population density in Fig 6–7 along each vertical column of lattice sites to obtain

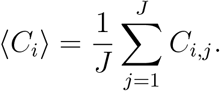

This quantity is further averaged by considering 100 identically prepared simulations of the discrete model to reduce fluctuations [19]. This procedure allows us to plot the time evolution of the average cell density as a function of the horizontal coordinate, as shown in Fig 8 [19]. Results in Fig 8a–b suggest that the evolution of the cell density profile is practically indistinguishable when we consider cells from the near–homogeneous cell line that is passaged without damage, as we might expect from the results in Fig 4a. In contrast, comparing results in Fig 8a with results in Fig 8c shows that we observe very different results when damage is included in the passaging process for the near–homogeneous cell line. When we consider the results in Fig 8d–f, for the heterogeneous cell line, we see that the evolution of the cell density profiles is very different for all three cases considered.

**Fig 8.**
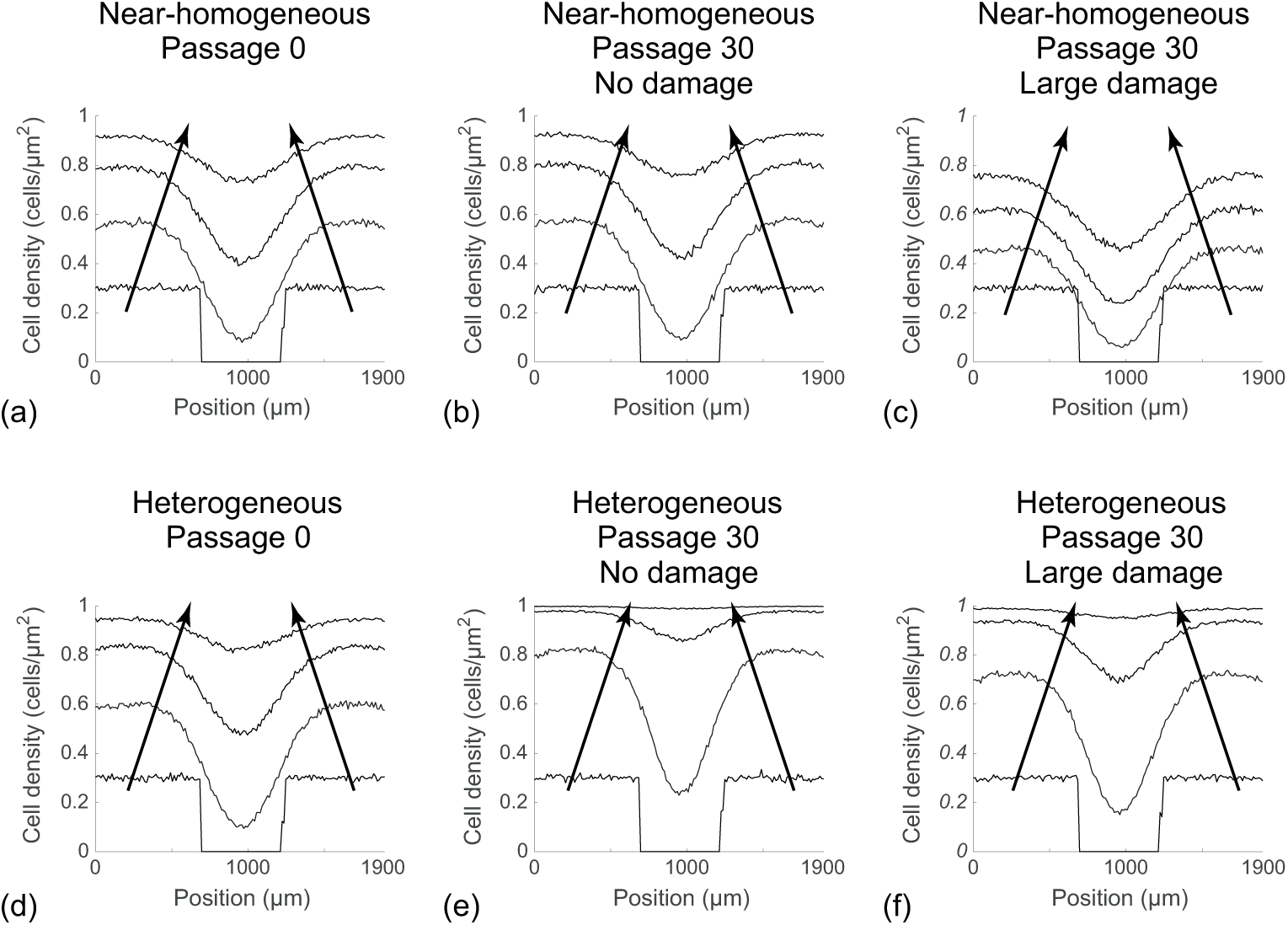
Averaged simulation data showing cell density profiles from the scratch assays. (a)–(c): Cell density profiles for a near–homogeneous cell line. (d)–(f): Cell density profiles for a heterogeneous cell line. In each subfigure cell density profiles are given at *t* = 0, 24, 48, 72 h, and the direction of increasing *t* is shown with the arrows. All simulation results are averaged across 100 identically prepared realisations of the discrete model, with Δ = 24 *μ*m, *τ* = 1/12 h, and *P*_*m*_ = 0.35, on a lattice of size 80 × 68. In each subfigure the initial *P*_*p*_ of individual cells is assigned by randomly selecting from the corresponding *in silico* data in Fig 2f, h, x, and Fig 3f, h, and x, respectively.

## Conclusion

Passaging of cell lines is an essential processes of growing cells in cell culture [3, 4]. The passaging process involves both chemical and mechanical disturbances which accumulatively change cell characteristics. Problems associated with high passage numbers, such as the change of cell proliferation, are widely acknowledged. However, the mechanisms are not well understood [10, 13, 18, 26, 29, 32, 34, 45]. Therefore, the aim of this work is to use a computational approach to provide insight into the putative mechanisms that could possibly lead to the problems.

In this work, we apply a lattice–based discrete model to investigate and quantify the impact of passaging cell lines. Although there are many properties of cells that are affected by the passaging process [6, 10, 13, 18, 31, 32, 34, 45], we choose to focus on how passaging affects the cell proliferation rate. In our model, when a cell proliferates, the daughter cells directly inherit the same proliferation rate as the mother cell. Furthermore, we also assume that during the passaging process, the cell proliferation rate is decreased by some passage–induced damage. For all results presented, we investigate the role of cell heterogeneity by comparing results where we begin the passaging process with a hear-homogeneous population of cells where *P*_*p*_ is almost constant, with a heterogeneous population of cells where *P*_*p*_ varies significantly among the population.

In the literature, previous experimental studies have reported apparently contradictory results where some studies suggest that the average proliferation rate of cells can increase at large passage number [26], whereas other studies suggest that the average proliferation rate of cells can decrease with passage number [10, 18, 32, 34]. We find that by varying the competition between passage-induced damage and cell heterogeneity, our relatively straightforward simulation model can predict each of these outcomes.

To study how passage number can affect *in vitro* experiments, we simulate a suite of scratch assays using various populations of cells that are harvested from our *in silico* passaging process. Our simulation results show that the passage number can lead to subtle changes in the evolution of the scratch assay and these changes might be very difficult to detect visually. We provide additional information about how the distribution of cells in a scratch assay might be influenced by passage number by performing a large number of realisations and examining the average cell density profiles. These average cell density profiles make it obvious that the passage number could affect the rate of scratch closure. This observation, together with the fact that cell passage number is often unreported in the experimental literature [2, 41], could explain why scratch assays are notoriously difficult to reproduce [12]. In addition, the results of cell culture growth and scratch assays indicate that even at the same passage number, the initial heterogeneity in cell proliferation can give rise to very differently behaving cell populations. Therefore, separating cell population without reporting the proliferative capacity can also affect the reproducibility of *in vitro* experiments. However, the proliferative capacity of cell lines can be difficult to measure experimentally, as most of the previous experiments only report the cell population evolution [18, 26], or the duration of the cell cycle [15].

There are several implications of this study that could be of interest to the experimental community. First, we suggest that the passage number of cell lines should always be reported. Second, there is a need for more experimental evidence about the impact of passaging on proliferation rates of various cell lines. For example, careful measurements of proliferation rates over a sequence of passage numbers would provide more insight into the variability of key cell properties in cell culture. This type of quantitative information would be invaluable for understanding reproducibility of standard *in vitro* experiments. Third, we acknowledge that our choices of the standard deviation, *σ*, to define the spread of the distribution of proliferation rates in the near–homogeneous and heterogeneous cell lines is rather theoretical. Recently, Haass et al. have devised new experimental methods that can be used to measure the durations of different phases in cell cycle for a range of melanoma cell lines [15]. This data could be used used to estimate the properties of the distribution of cell proliferation rates, such as the mean and standard deviation of the distribution of proliferation rates. Therefore, we suggest that similar experiments could be performed to generate proliferation rate distribution over various passage numbers for a range of different cell lines of interest. This data could then be directly integrated within our *in silico* models to examine the interplay between the degree of heterogeneity and passage–induced damage.

There are also several implications of this study that are of interest to the applied mathematics and mathematical biology communities. First, here we focus on the case where there is heterogeneity in the rate at which individual cells proliferate in the population but, we treat the motility rate as a constant. This is because most previous experimental studies have reported differences in the rate of proliferation as a function of passage number rather than differences in the rate of migration [18, 26, 34]. However, heterogeneity in cell migration rate can also affect the reproducibility of *in vitro* experiments [32], especially scratch assays in which cell migration plays a key role in wound closure [20]. An interesting extension of our present study would involve dealing with both variability in the motility rate and the proliferation rate [30]. Secondly, in our work we make the most straightforward assumption that daughter cells inherit *P*_*p*_ directly from the mother cell. It might be more plausible to introduce some stochasticity in the inheritance process, and it might also be plausible to incorporate some kind of ageing process where the proliferation depends on the age structure of the population [5, 32]. We have chosen not to include these additional details as we wish to present a simpler model that is capable of illustrating a proof–of–principle concept rather than capturing every possible feature of the underlying biology. Finally, another extension of this work would be to consider the derivation of an accurate mean–field approximation that could be used to describe the evolution of the cell density profiles in Fig 7. This is a challenging task because all previous derivations of such mean–field partial differential equations involve populations of cells with constant rates [7, 19, 27, 35, 36, 40], whereas we are dealing with a more realistic heterogeneous population of cells.

## Acknowledgments

This work is supported by the Australian Research Council (DP140100249, DP170100474), and we appreciate the helpful comments from the two reviewers.

